# Quantifying concordant genetic effects of *de novo* mutations on multiple disorders

**DOI:** 10.1101/2021.06.13.448234

**Authors:** Hanmin Guo, Lin Hou, Yu Shi, Sheng Chih Jin, Xue Zeng, Boyang Li, Richard P. Lifton, Martina Brueckner, Hongyu Zhao, Qiongshi Lu

## Abstract

Exome sequencing on tens of thousands of parent-proband trios has identified numerous deleterious *de novo* mutations (DNMs) and implicated risk genes for many disorders. Recent studies have suggested shared genes and pathways are enriched for DNMs across multiple disorders. However, existing analytic strategies only focus on genes that reach statistical significance for multiple disorders and require large trio samples in each study. As a result, these methods are not able to characterize the full landscape of genetic sharing due to polygenicity and incomplete penetrance. In this work, we introduce EncoreDNM, a novel statistical framework to quantify shared genetic effects between two disorders characterized by concordant enrichment of DNMs in the exome. EncoreDNM makes use of exome-wide, summary-level DNM data, including genes that do not reach statistical significance in single-disorder analysis, to evaluate the overall and annotation-partitioned genetic sharing between two disorders. Applying EncoreDNM to DNM data of nine disorders, we identified abundant pairwise enrichment correlations, especially in genes intolerant to pathogenic mutations and genes highly expressed in fetal tissues. These results suggest that EncoreDNM improves current analytic approaches and may have broad applications in DNM studies.

## Introduction

*De novo* mutations (DNMs) can be highly deleterious and provide important insights into the genetic cause for disease^1^. As the cost of sequencing continues to drop, whole-exome sequencing (WES) studies conducted on tens of thousands of family trios have pinpointed numerous risk genes for a variety of disorders^2–4^. In addition, accumulating evidence suggests that risk genes enriched for pathogenic DNMs may be shared by multiple disorders^5–9^. These shared genes could reveal biological pathways that play prominent roles in disease etiology and shed light on clinically heterogeneous yet genetically related diseases^7–9^.

Most efforts to identify shared risk genes directly compare genes that are significantly associated with each disorder^10,11^. There have been some successes with this approach in identifying shared genes and pathways (e.g., chromatin modifiers) underlying developmental disorder (DD), autism spectrum disorder (ASD), and congenital heart disease (CHD), thanks to the large trio samples in these studies^3,4,12^, whereas findings in smaller studies remain suggestive^13,14^. Even in the largest studies to date, statistical power remains moderate for risk genes with weaker effects^3,15^. It is estimated that more than 1,000 genes associated with DD remain undetected^3^. Therefore, analytic approaches that only account for top significant genes cannot capture the full landscape of genetic sharing in multiple disorders. Recently, a Bayesian framework was proposed to jointly analyze DNM data of two diseases and improve risk gene mapping^9^. Although some parameters in this framework can quantify shared genetics between diseases, the statistical property of these parameter estimates have not been studied. There is a pressing need for powerful, robust, and interpretable methods that quantify concordant DNM association patterns for multiple disorders using exome-wide DNM counts.

Recent advances in estimating genetic correlations using summary data from genome-wide association studies (GWAS) may provide a blueprint for approaching this problem in DNM research^16^. Modeling “omnigenic” associations as independent random effects, linear mixed-effects models leverage genome-wide association profiles to quantify the correlation between additive genetic components of multiple complex traits^17–20^. These methods have identified ubiquitous genetic correlations across many human traits and revealed significant and robust genetic correlations that could not be inferred from significant GWAS associations alone^21–24^.

Here, we introduce EncoreDNM (**En**richment **cor**relation **e**stimator for ***D**e **N**ovo* **M**utations), a novel statistical framework that leverages exome-wide DNM counts, including genes that do not reach exome-wide statistical significance in single-disorder analysis, to estimate concordant DNM associations between disorders. EncoreDNM uses a generalized linear mixed-effects model to quantify the occurrence of DNMs while accounting for *de novo* mutability of each gene and technical inconsistencies between studies. We demonstrate the performance of EncoreDNM through extensive simulations and analyses of DNM data of nine disorders.

## Results

### Method overview

DNM counts in the exome deviate from the null (i.e., expected counts based on *de novo* mutability) when mutations play a role in disease etiology. Disease risk genes will show enrichment for deleterious DNMs in probands and non-risk genes may be slightly depleted for DNM counts. Our goal is to estimate the correlation of such deviation between two disorders, which we refer to as the DNM enrichment correlation. More specifically, we use a pair of mixed-effects Poisson regression models to quantify the occurrence of DNMs in two studies.

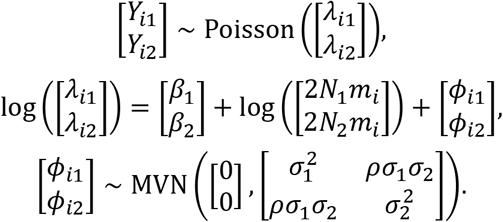

Here, *Y*_*i*1_, *Y*_*i*2_ are the DNM counts for the *i*-th gene and *N*_1_, *N*_2_ are the number of parent-proband trios in two studies, respectively. The log Poisson rates of DNM occurrence are decomposed into three components: the elevation component, the background component, and the deviation component. The elevation component *β_k_* (*k* = 1,2) is a fixed effect term adjusting for systematic, exome-wide bias in DNM counts. One example of such bias is the batch effect caused by different sequencing and variant calling pipelines in two studies. The elevation parameter *β_k_* tends to be larger when DNMs are over-called with higher sensitivity and smaller when DNMs are under-called with higher specificity^25^. The background component log(2*N_k_m_i_*) is a gene-specific fixed effect that reflects the expected mutation counts determined by the genomic sequence context under the null^26^. *m_i_* is the *de novo* mutability for the *i*-th gene, and 2*N*_1_*m_i_* and 2*N*_2_*m_i_* are the expected DNM counts in the *i*-th gene under the null in two studies. The deviation component *ϕ_ik_* is a gene-specific random effect that quantifies the deviation of DNM profile from what is expected under the null. *ϕ*_*i*1_ and *ϕ*_*i*2_ follow a multivariate normal distribution with dispersion parameters *σ*_1_ and *σ*_2_ and a correlation *ρ*. DNM enrichment correlation is denoted by *ρ* and is our main parameter of interest. It quantifies the concordance of DNM burden in two disorders.

Parameters in this model can be estimated using a Monte Carlo maximum likelihood estimation (MLE) procedure. Standard errors of the estimates are obtained through inversion of the observed Fisher information matrix. In practice, we use annotated DNM data as input and fit mixed-effects Poisson models for each variant class separately: loss of function (LoF), deleterious missense (Dmis, defined as MetaSVM-deleterious), tolerable missense (Tmis, defined as MetaSVM-tolerable), and synonymous (**Figure 1**). More details about model setup and parameter estimation are discussed in **Methods**.

**Figure 1.**
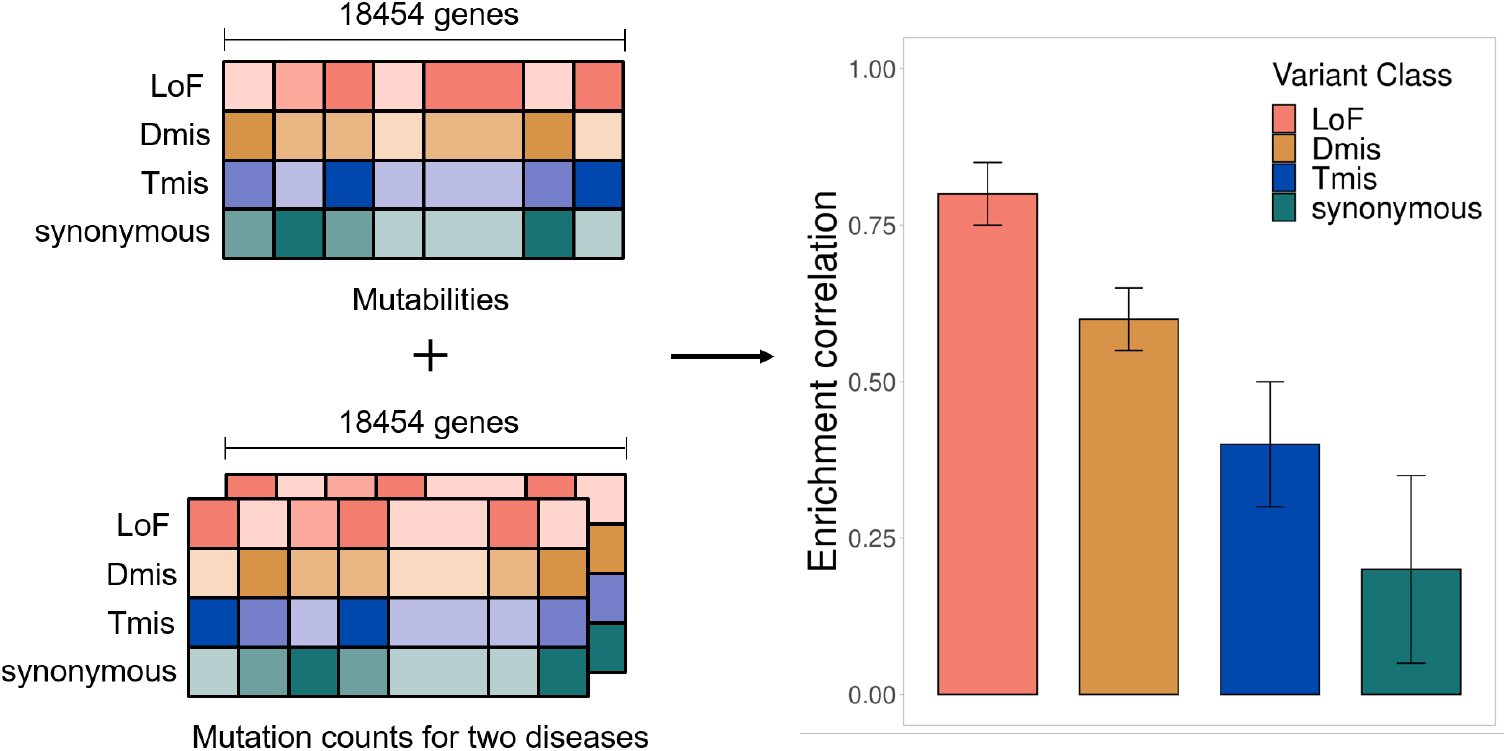
EncoreDNM workflow. The inputs of EncoreDNM are *de novo* mutability of each gene and exome-wide, annotated DNM counts from two studies. We fit a mixed-effects Poisson model to estimate the DNM enrichment correlation between two disorders for each variant class.

### Simulation results

We conducted simulations to assess the parameter estimation performance of EncoreDNM in various settings. We focused on two variant classes, i.e., Tmis and LoF variants, since they have the highest and lowest median mutabilities in the exome. We used EncoreDNM to estimate the elevation parameter *β*, dispersion parameter *σ*, and enrichment correlation *ρ* (**Methods**). Under various parameter settings, EncoreDNM always provided unbiased estimation of the parameters (**Figure 2** and **Supplementary Figures 1-2**). Furthermore, the 95% Wald confidence intervals achieved coverage rates close to 95% under all simulation settings, demonstrating the effectiveness of EncoreDNM to provide accurate statistical inference.

**Figure 2.**
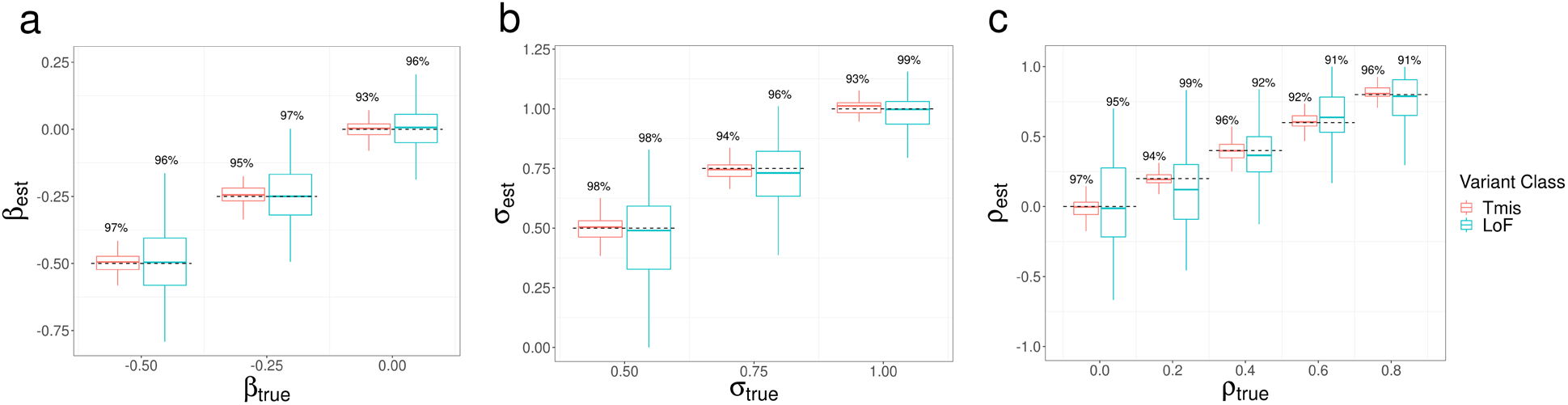
Parameter estimation results of EncoreDNM. (a) Boxplot of *β* estimates in single-trait analysis with *σ* fixed at 0.75. (b) Boxplot of *σ* estimates in single-trait analysis with *β* fixed at −0.25. (c) Boxplot of *ρ* estimates in cross-trait analysis with *β* and *σ* fixed at −0.25 and 0.75. True parameter values are marked by dashed lines. The number above each box represents the coverage rate of 95% Wald confidence intervals. Each simulation setting was repeated 100 times.

Next, we compared the performance of EncoreDNM with mTADA^9^, a Bayesian framework that could estimate the proportion of shared risk genes for two disorders. First, we simulated DNM data under the mixed-effects Poisson model. We evaluated two methods across a range of combinations of elevation parameter, dispersion parameter, and sample size for two disorders. The type-I error rates for our method were well-calibrated in all parameter settings, but mTADA produced false positive findings when the observed DNM counts were relatively small (e.g., due to reduced elevation or dispersion parameters or a lower sample size; **Figure 3a**). We also assessed the statistical power of two approaches under a baseline setting where type-I errors for both methods were controlled. As enrichment correlation increased, EncoreDNM achieved universally greater statistical power compared to mTADA (**Figure 3b**).

**Figure 3.**
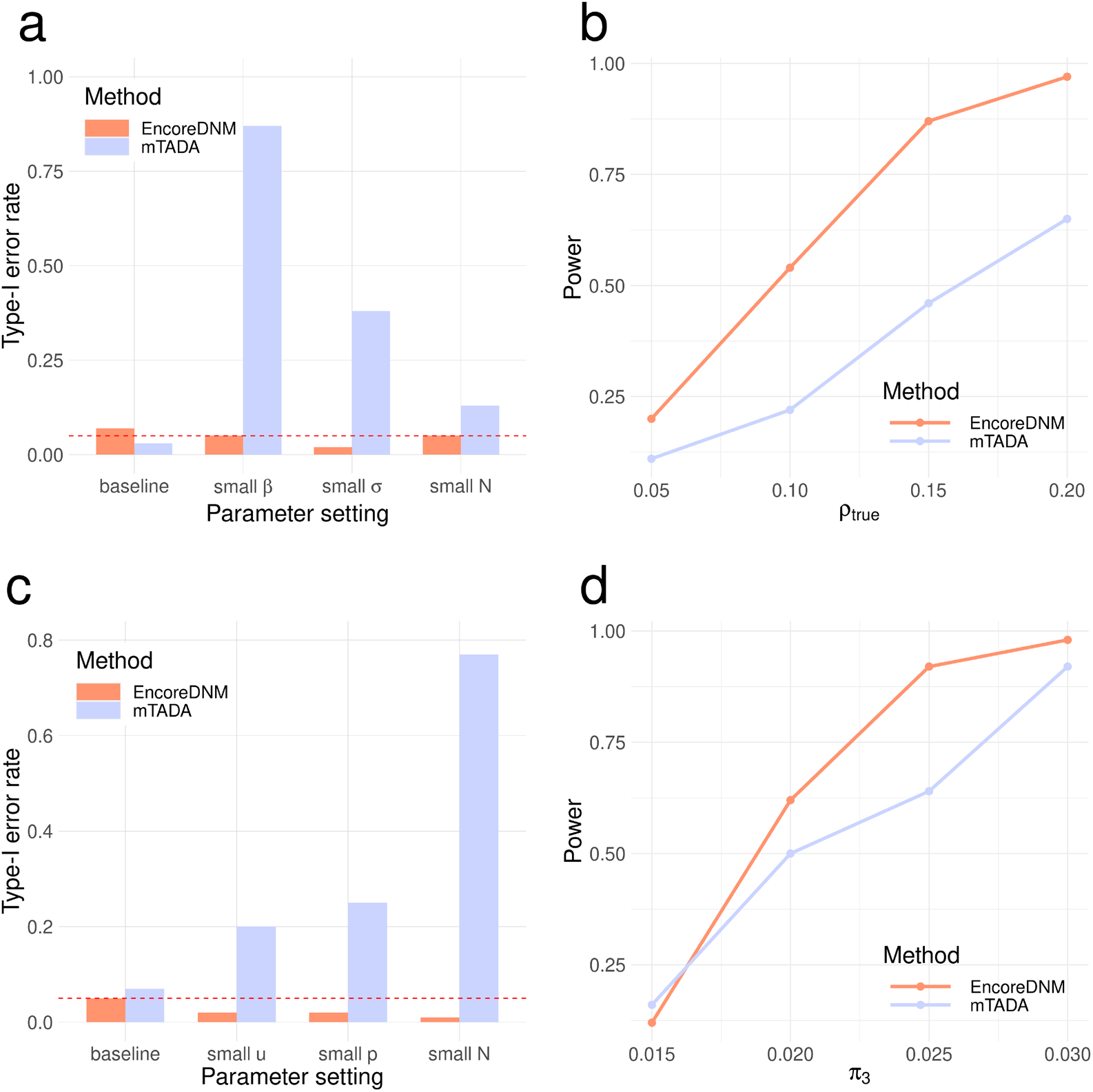
Comparison of EncoreDNM and mTADA. (a) Type-I error rates under a mixed-effects Poisson regression model. (*β*, *σ*, *N*) were fixed at (−0.25, 0.75, 5000) under the baseline setting, (−1, 0.75, 5000) under a setting with small *β*, (−0.25, 0.5, 5000) under a setting with small *σ*, and (−0.25, 0.75, 1000) under a setting with small N for two disorders. (b) Statistical power of two methods under a mixed-effects Poisson regression model as the enrichment correlation increases. Parameters (*β*, *σ*, *N*) were fixed at (−0.25, 0.75, 5000) for both disorders. (c) Type-I error rates under a multinomial model. (*u*, *p*, *N*, *π^s^*) were fixed at (0.95, 0.25, 5000, 0.1) under the baseline setting, (0.75, 0.25, 5000, 0.1) under a setting with small u (i.e., reduced total DNM counts), (0.95, 0.15, 5000, 0.1) under a setting with small p (i.e., fewer probands explained by DNMs), and (0.95, 0.25, 1000, 0.1) under a setting with lower sample size. (d) Statistical power under a multinomial model with varying proportion of shared causal genes. Parameters (*u*, *p*, *N*, *π^s^*) were fixed at (0.95, 0.25, 5000, 0.1) for both disorders. Each simulation setting was repeated 100 times.

To ensure a fair comparison, we also considered a mis-specified model setting where we randomly distributed the total DNM counts for each disorder into all genes with an enrichment in causal genes (**Methods**). EncoreDNM showed well-controlled type-I error across all simulation settings, whereas severe type-I error inflation arose for mTADA when the total mutation count, the proportion of probands that can be explained by DNMs, or the sample size were small (**Figure 3c**). Furthermore, we compared the statistical power of two methods under this model in a baseline setting where type-I error was controlled. EncoreDNM showed higher statistical power compared to mTADA as the fraction of shared causal genes increased (**Figure 3d**).

### Pervasive enrichment correlation of damaging DNMs among developmental disorders

We applied EncoreDNM to DNM data of nine disorders (**Supplementary Table 1**; **Methods**): DD (n=31,058; number of trios)^3^, ASD (n=6,430)^4^, schizophrenia (SCZ; n=2,772)^15^, CHD (n=2,645)^12^, intellectual disability (ID; n=820)^2^, Tourette disorder (TD; n=484)^27^, epileptic encephalopathies (EP; n=264)^13^, cerebral palsy (CP; n=250)^14^, and congenital hydrocephalus (CH; n=232)^28^. In addition, we also included 1,789 trios comprising healthy parents and unaffected siblings of ASD probands as controls^29^.

We first performed single-trait analysis for each disorder. The estimated elevation parameters (i.e., *β*) were negative for almost all disorders and variant classes (**Figure 4a**), with LoF variants showing particularly lower parameter estimates. This may be explained by more stringent quality control in LoF variant calling^12^ and potential survival bias^30^. It is also consistent with a depletion of LoF DNMs in healthy control trios^7^. The dispersion parameter estimates (i.e., *σ*) were higher for LoF variants than other variant classes (**Figure 4b**), which is consistent with our expectation that LoF variants have stronger effects on disease risk and should show a larger deviation from the null mutation rate in disease probands. We compared the goodness of fit of our proposed mixed-effects Poisson model to a simpler fixed-effects model without the deviation component (**Methods**). The expected distribution of recurrent DNM counts showed substantial and statistically significant improvement under the mixed-effects Poisson model (**Figures 4c-f** and **Supplementary Figure 3**).

**Figure 4.**
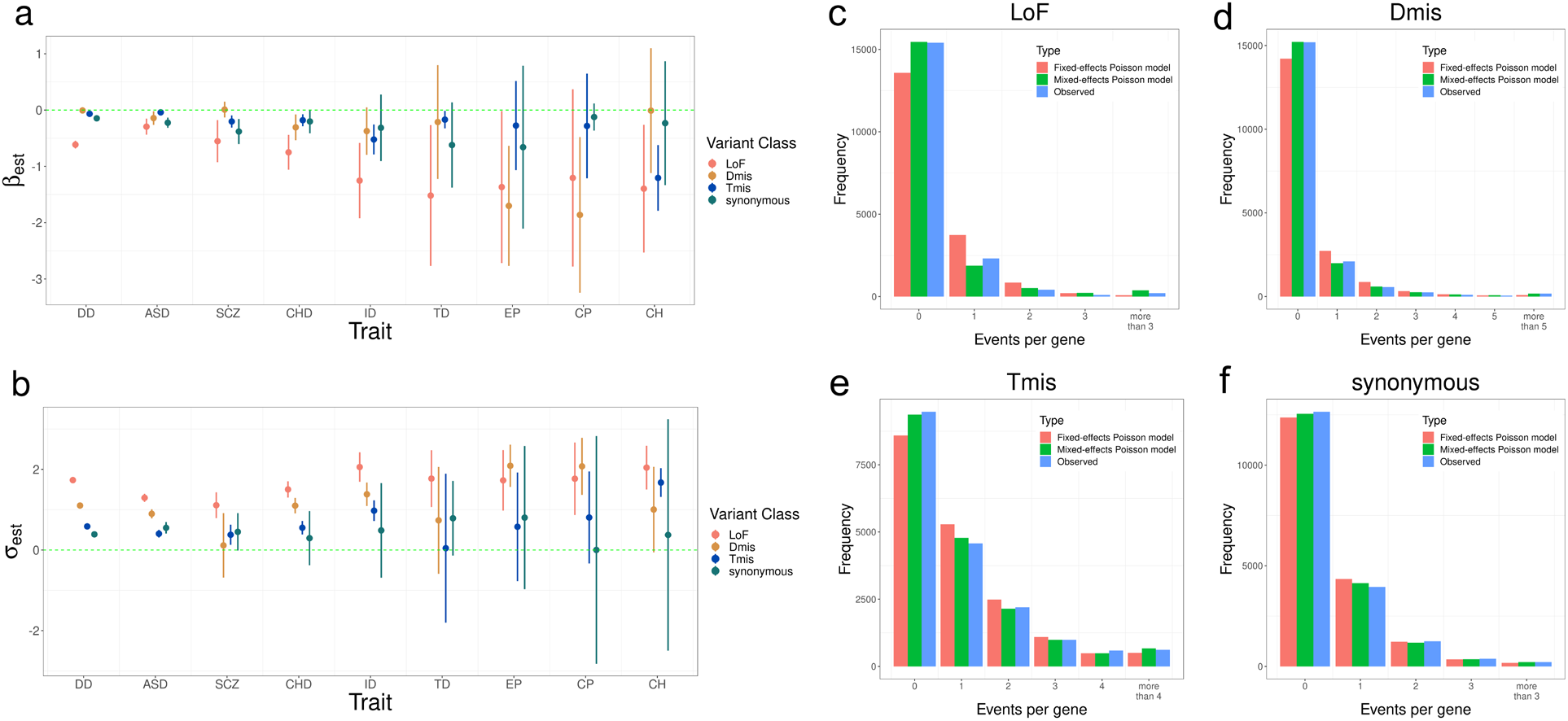
Model fitting results for nine disorders. (a, b) Estimation results of *β* and *σ* for nine disorders and four variant classes. Error bars represent 1.96*standard errors. (c-f) Distribution of DNM events per gene in four variant classes for DD. Red and green bars represent the expected frequency of genes under the fixed-effects and mixed-effects Poisson regression models, respectively. Blue bars represent the observed frequency of genes.

Next, we estimated pairwise DNM enrichment correlations for 9 disorders. In total, we identified 25 pairs of disorders with significant correlations at a false discovery rate (FDR) cutoff of 0.05 (**Figure 5** and **Supplementary Figure 4**), including 12 significant correlations for LoF variants, 7 for Dmis variants, 5 for Tmis variants, and only 1 significant correlation for synonymous variants. Notably, all significant correlations are positive (**Supplementary Table 2**). No significant correlation was identified between any disorder and healthy controls (**Supplementary Figure 5**). The number of identified significant correlations for each disorder was proportional to the sample size in each study (Spearman correlation = 0.70) with controls being a notable outlier (**Supplementary Figure 6**).

**Figure 5.**
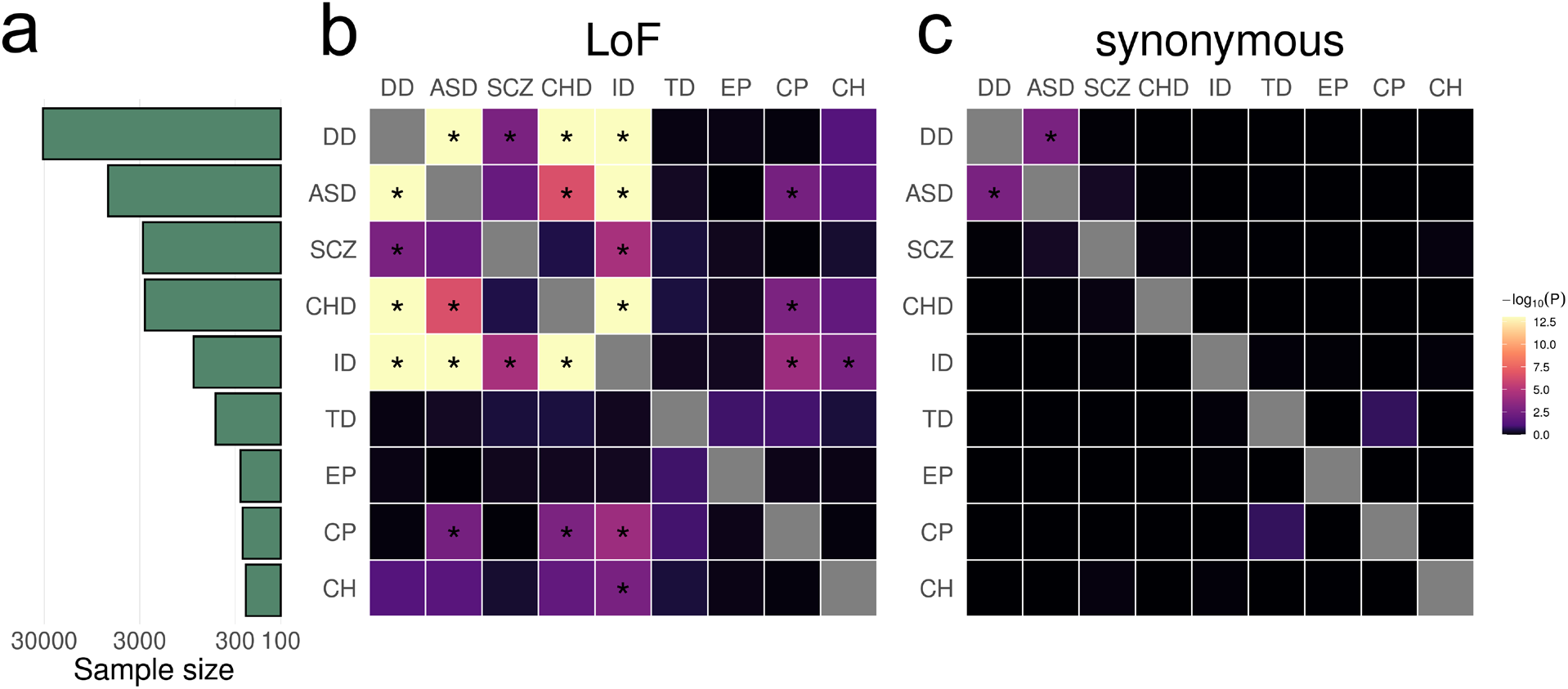
EncoreDNM identifies pervasive enrichment correlations of damaging DNMs among nine disorders. (a) shows sample size (i.e., number of trios) for each disease. X-axis denotes sample size on the log scale. (b, c) Heatmap of enrichment correlations for LoF and synonymous DNMs among nine disorders. Significant correlations (FDR<0.05) are marked by asterisks. Results with −log_10_ *P* >13 are truncated to 13 for visualization purpose.

We identified highly concordant and significant LoF DNM enrichment among DD, ASD, ID, and CHD, which is consistent with previous reports^8–10^,^31^. SCZ shows highly significant LoF correlations with DD and ID (p=2.0e-3 and 3.7e-5), hints at a correlation with ASD (p=0.012), but does not correlate strongly with CHD. The positive enrichment correlation between ASD and CP in LoF variants (*ρ*=0.81, p=3.3e-3) is consistent with their co-occurrence^32^. The high enrichment correlation between ID and CP in LoF variants (*ρ* =0.68, p=1.0e-4) is consistent with the associations between ID and motor or non-motor abnormalities caused by CP^33^. A previous study also suggested significant genetic sharing of ID and CP by overlapping genes harboring rare damaging variants^14^. Here, we obtained consistent results after accounting for *de novo* mutabilities and potential confounding bias.

Some significant correlations identified in our analysis are consistent with phenotypic associations in epidemiological studies, but have not been reported using genetic data to the extent of our knowledge. For example, the LoF enrichment correlation between CHD and CP (*ρ*=0.88, p=1.7e-3) is consistent with findings that reduced supply of oxygenated blood in fetal brain due to cardiac malformations may be a risk factor for CP^34^. The enrichment correlation between ID and CH in LoF variants (*ρ*=0.63, p=2.4e-3) is consistent with lower intellectual performance in a proportion of children with CH^35^.

Genes showing pathogenic DNMs in multiple disorders may shed light on the mechanisms underlying enrichment correlations (**Supplementary Table 3**). We identified five genes (i.e. *CTNNB1*, *NBEA*, *POGZ*, *SPRED2*, and *KMT2C*) with LoF DNMs in five different disorders and 21 genes had LoF DNMs in four disorders (**Supplementary Table 4**). These 26 genes with LoF variants in at least four disorders were significantly enriched for 63 gene ontology (GO) terms with FDR<0.05 (**Supplementary Table 5**). Chromatin organization (p=7.8e-11), nucleoplasm (p=2.8e-10), chromosome organization (p=6.8e-10), histone methyltransferase complex (p=1.4e-9), and positive regulation of gene expression (p=2.2e-9) were the most significantly enriched GO terms. One notable example consistently included in these gene sets is *CTNNB1* (**Supplementary Figure 7**). It encodes *β*-catenin, is one of the only two genes reaching genome-wide significance in a recent WES study for CP^14^, and also harbors multiple LoF variants in DD, ID, ASD, and CHD. It is a fundamental component of the canonical Wnt signaling pathway which is known to confer genetic risk for ASD^36^. We also identified 157 recurrent LoF mutations in 45 genes (**Supplementary Table 6**). Most of these recurrent mutations were identified in DD due to its large sample size, but one mutation was identified in joint comparison of other disorders. *FBXO11*, encoding the F-box only protein 31, shows two recurrent p.Ser831fs LoF variants in ASD and CH (**Supplementary Figure 8**; p=1.9e-3; **Methods**). The F-box protein constitutes a substrate-recognition component of the SCF (SKP1-cullin-F-box) complex, an E3-ubiquitin ligase complex responsible for ubiquitination and proteasomal degradation^37^. DNMs in *FBXO11* have been previously implicated in severe ID individuals with autistic behavior problem^38^ and neurodevelopmental disorder^39^.

For comparison, we also applied mTADA to the same nine disorders and control trios. In total, mTADA identified 117 disorder pairs with significant genetic sharings at an FDR cutoff of 0.05 (**Supplementary Table 7** and **Supplementary Figure 9**). Notably, we identified significant synonymous DNM correlations for all 36 disorder pairs and between all disorders and healthy controls (**Supplementary Figure 10**). These results are consistent with the simulation results and suggest a substantially inflated false positive rate in mTADA.

Further, we applied cross-trait linkage disequilibrium (LD) score regression^18^ to five of the nine disorders with publicly available GWAS summary statistics (**Supplementary Table 8**): ASD (n=46,350)^40^, SCZ (n=161,405)^41^, cognitive performance (used as a proxy for ID; n=257,841)^42^, TD (n=14,307)^43^, and epilepsy (n=44,889)^44^. In total, we identified 6 trait pairs with significant genetic correlations at an FDR cutoff of 0.05 (**Supplementary Table 9**), suggesting consistent findings made from GWAS and DNM data (Spearman correlation = 0.70; **Supplementary Figure 11**).

### Partitioning DNM enrichment correlation by gene set

To gain biological insights into the shared genetic architecture of nine disorders, we repeated EncoreDNM correlation analysis in several gene sets. First, we defined genes with high/low probability of intolerance to LoF variants using pLI scores^45^, and identified genes with high/low brain expression (HBE/LBE)^46^ (**Methods**; **Supplementary Table 10**). We identified 11 and 12 disorder pairs showing significant enrichment correlations for LoF DNMs in high-pLI genes and HBE genes, respectively (**Figure 6a-b**). We observed fewer significant correlations for Dmis and Tmis variants in these gene sets (**Supplementary Figures 12-13**). All identified significant correlations were positive (**Supplementary Tables 11-12**). No significant correlations were identified for synonymous variants (**Supplementary Figures 12-13**) or between disorders and controls (**Supplementary Figures 14-15**).

**Figure 6.**
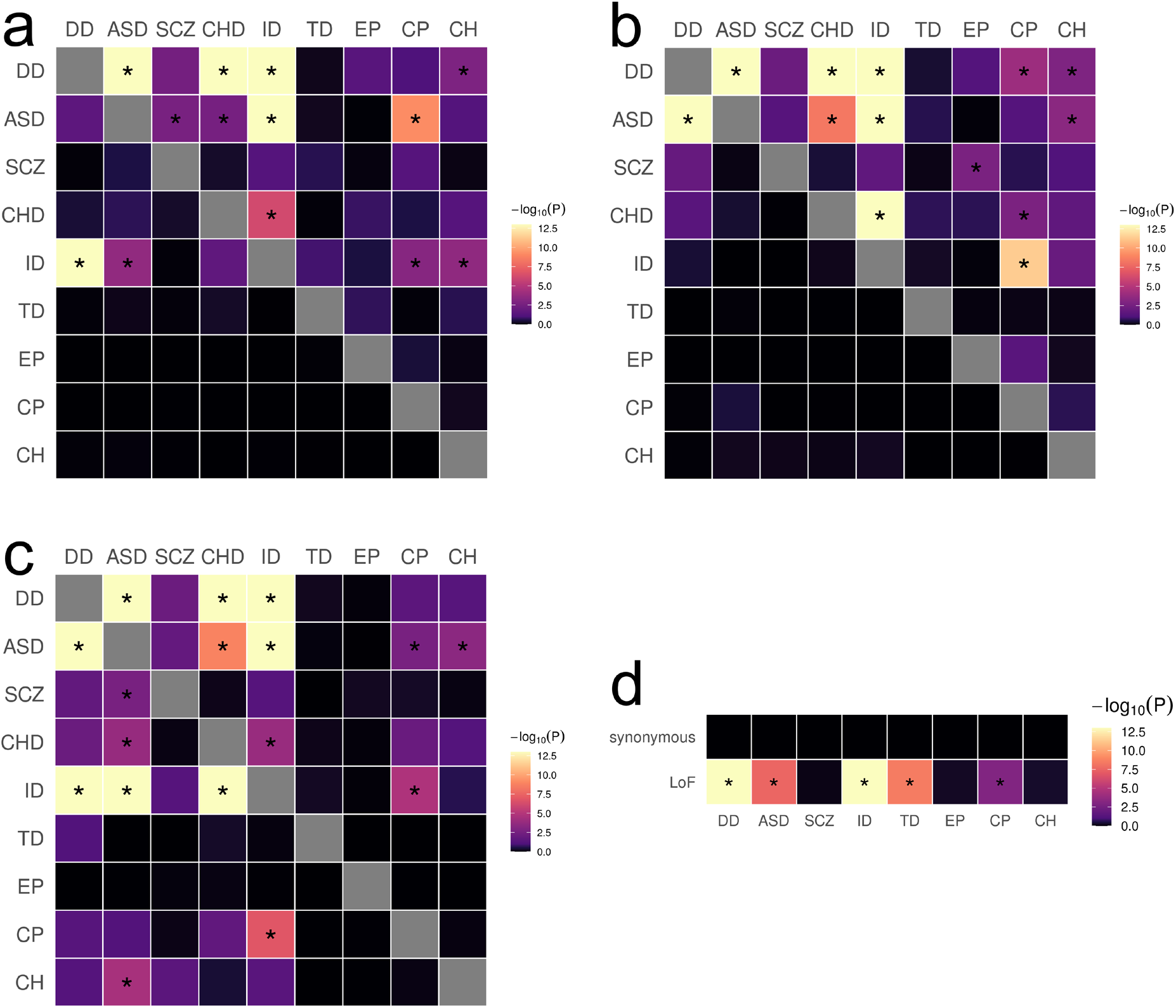
DNM enrichment correlations in disease-relevant gene sets. (a) Enrichment correlations in High-pLI genes (upper triangle) and Low-pLI genes (lower triangle) for LoF variants. (b) Enrichment correlations in HBE genes (upper triangle) and LBE genes (lower triangle) for LoF variants. (c) Enrichment correlations in HHE genes (upper triangle) and LHE genes (lower triangle) for LoF variants. (d) Enrichment correlations in CHD-related pathways for LoF and synonymous variants. Significant correlations (FDR<0.05) are marked by asterisks. Results with −log_10_ P > 13 are truncated to 13 for visualization purpose.

We observed a clear enrichment of significant correlations in disease-relevant gene sets. Overall, high-pLI genes showed substantially stronger correlations across disorders than genes with low pLI (one-sided Kolmogorov-Smirnov test; p=2.3e-6). Similarly, enrichment correlations were stronger in HBE genes than in LBE genes (p=8.8e-7). Among the 11 disorder pairs showing significant enrichment correlations in high-pLI genes, two pairs, i.e., ASD-SCZ (*ρ*=0.68, p=2.4e-3) and DD-CH (*ρ*=0.43, p=1.5e-3), were not identified in the exome-wide analysis. We also identified four novel disorder pairs with significant correlations in HBE genes, including DD-CP (*ρ* =0.80, p=9.5e-5), DD-CH (*ρ* =0.67, p=1.4e-3), ASD-CH (*ρ* =0.82, p=4.7e-4), and SCZ-EP (*ρ*=0.66, p=2.0e-3). These novel enrichment correlations are consistent with known comorbidities between these disorders^47,48^ and findings based on significant risk genes^8,28,49,50^.

Furthermore, we estimated DNM enrichment correlations in genes with high/low expression in mouse developing heart (HHE/LHE)^7^ (**Methods**; **Supplementary Table 10**). We identified 9 significant enrichment correlations for LoF variants in HHE genes (**Figure 6c**). Strength of enrichment correlations did not show a significant difference between HHE and LHE genes (p=0.846), possibly due to a lack of cardiac disorders in our analysis. Finally, we estimated enrichment correlations between CHD and other disorders in known pathways for CHD^51^ (**Methods**; **Supplementary Table 10**). We identified 5 significant correlations for LoF variants (**Figure 6d**), including a novel correlation between CHD and TD (*ρ*=0.93, p=3.3e-9). Of note, arrhythmia caused by CHD is a known risk factor for TD^52^. In these analyses, all significant enrichment correlations were positive (**Supplementary Tables 13-14**) and other variant classes showed generally weaker correlations than LoF variants (**Supplementary Figures 16-17**). We did not observe significant correlations in these gene sets between disorders and controls (**Supplementary Figures 18-19**).

## Discussion

In this paper, we introduced EncoreDNM, a novel statistical framework to quantify correlated DNM enrichment between two disorders. Through extensive simulations and analyses of DNM data for nine disorders, we demonstrated that our proposed mixed-effects Poisson regression model provides unbiased parameter estimates, shows well-controlled type-I error, and is robust to exome-wide technical biases. Leveraging exome-wide DNM counts and genomic context-based mutability data, EncoreDNM achieves superior fit for real DNM datasets compared to simpler models and provides statistically powerful and computationally efficient estimation of DNM enrichment correlation. Further, EncoreDNM can quantify concordant genetic effects for user-defined variant classes within pre-specified gene sets, thus is suitable for exploring diverse types of hypotheses and can provide crucial biological insights into the shared genetic etiology in multiple disorders.

Multi-trait analyses of GWAS data have revealed shared genetic architecture among many neuropsychiatric traits^22,53,54^. These findings have led to the identification of pleiotropic variants, genes, and hub genomic regions underlying many traits and have revealed multiple psychopathological factors jointly affecting human neurological phenotypes^55,56^. Although emerging evidence suggests that causal DNMs underlying several disorders with well-powered studies (e.g., CHD and neurodevelopmental disorders^7^) may be shared, our understanding of the extent and the mechanism underlying such sharing remains incomplete. Applied to DNM data for nine disorders, EncoreDNM identified pervasive enrichment correlations of DNMs. We observed particularly strong correlations in pathogenic variant classes (e.g., LoF and Dmis variants) and disease-relevant genes (e.g., genes with high pLI and genes highly expressed in relevant tissues). Genes underlying these correlations were significantly enriched in pathways involved in chromatin organization and modification and gene expression regulation. The DNM correlations were substantially attenuated in genes with lower expression and genes with frequent occurrences of LoF variants in the population. A similar attenuation was observed in less pathogenic variant classes (e.g., synonymous variants). Further, no significant correlations were identified between any disorder and healthy controls. These results lay the groundwork for future investigations of pleiotropic mechanisms of DNMs.

Our study has some limitations. First, a main goal in DNM research is to identify disease risk genes. EncoreDNM leverages exome-wide DNM counts to quantify shared genetic basis in multiple disorders but does not improve the analysis of gene-disease associations. Second, EncoreDNM assumes probands from different input studies to be independent. In rare cases when two studies have overlapping proband samples, enrichment correlation estimates may be inflated and must be interpreted with caution. Finally, genetic correlation methods based on GWAS summary data provided key motivations for the mixed-effects Poisson regression model in our study. Built upon genetic correlations, a plethora of methods have been developed in the GWAS literature to jointly model more than two GWAS^57^, identify and quantify common factors underlying multiple traits^58,59^, estimate causal effects among different traits^60^, and identify pleiotropic genomic regions through hypothesis-free scans^23^. Future directions of EncoreDNM include using enrichment correlation to improve gene discovery, learning the directional effects and the causal structure underlying multiple disorders, and dynamically searching for gene sets and annotation classes with shared genetic effects without pre-specifying the hypothesis.

Taken together, we provide a new analytic approach to an important problem in DNM studies. We believe EncoreDNM improves the statistical rigor in multi-disorder DNM modeling and opens up many interesting future directions in both method development and follow-up analyses in WES studies. As trio sample size in WES studies continues to grow, EncoreDNM will have broad applications and can greatly benefit DNM research.

## Methods

### Statistical Model

For a single study, we assume that DNM counts in a given variant class (e.g., synonymous variants) follow a mixed-effects Poisson model:

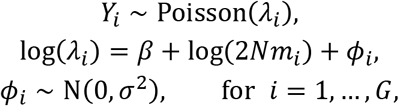

where *Y_i_* is the DNM count in the *i*-th gene, *N* is the number of trios, *m_i_*, is the *de novo* mutabilityfor the *i*-th gene (i.e., mutation rate per chromosome per generation) which is known *a priori*^26^ (**Supplementary Table 15**), and *G* is the total number of genes in the study. The elevation parameter *β* quantifies the global elevation of mutation rate compared to mutability estimates based on genomic sequence alone. Gene-specific deviation from expected DNM rate is quantified by random effect *ϕ_i_* with a dispersion parameter *σ*. Here, the *ϕ_i_* are assumed to be independent across different genes, in which case the observed DNM counts of different genes are independent.

Next, we describe how we expand this model to quantify the shared genetics of two disorders. We assume DNM counts in a given variant class for two diseases follow:

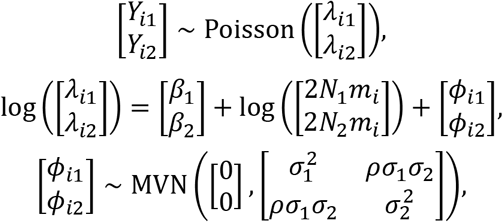

where *Y*_*i*1_, *Y*_*i*2_ are the DNM counts for the *i*-th gene and *N*_1_, *N*_2_ are the trio sizes in two studies, respectively. Similar to the single-trait model, is the mutability for the *i*-th gene. *β*_1_, *β*_2_ are the elevation parameters, and *ϕ*_*i*1_, *ϕ*_*i*2_ are the gene-specific random effects with dispersion parameters *σ*_1_, *σ*_2_, for two disorders respectively. *ρ* is the enrichment correlation which quantifies the concordance of the gene-specific DNM burden between two disorders. Here, *β*_1_, *β*_2_, *σ*_1_, *σ*_2_, *ρ* are unknown parameters. The gene specific effects for two disorders are assumed to be independent for different genes. We also assume that there is no shared sample for two disorders, in which case *Y*_*i*1_ is independent with *Y*_i_2 given 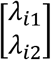.

### Parameter estimation

We implement an MLE procedure to estimate unknown parameters. For single-trait analysis, the log-likelihood function can be expressed as follows:

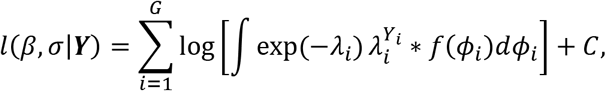

where ***Y*** = [*Y*_1_,… *Y_G_*]^T^, *λ_i_* = 2*Nm_i_* exp(*β* + *ϕ_i_*), 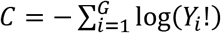, and 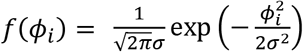. Note that there is no closed form for the integral in the log-likelihood function. Therefore, we use Monte Carlo integration to evaluate the log-likelihood function. Let *ϕ_ij_* = *σξ_ij_*, where the *ξ_ij_* are independently and identically distributed random variables following a standard normal distribution. We have

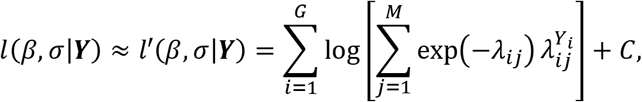

where *λ_ij_* = 2*Nm_i_* exp(*β* + *σξ_ij_*), and *M* is the Monte Carlo sample size which is set to be 1000. Then, we could obtain the MLE of *β*, *σ* through maximization of *l’*(*β*, *σ*|***Y***). We obtain the standard error of the MLE through inversion of the observed Fisher information matrix.

The estimation procedure can be generalized to multi-trait analysis. Log-likelihood function can be expressed as follows:

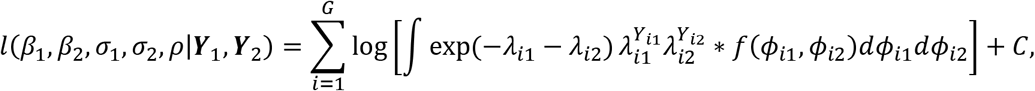

where ***Y***_1_ = [*Y*_11_,…, *Y*_*G*1_]^T^, ***Y***_2_ = [*Y*_12_,…, *Y*_*G*2_]^T^, *λ*_*i*1_ = 2*N*_1_*m_i_* exp(*β*_1_ + *ρ*_*i*1_), *λ*_*i*2_ = 2*N*_2_*m_i_* exp(*β*_2_ + *ϕ*_*i*2_), 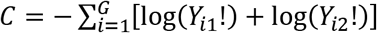, and 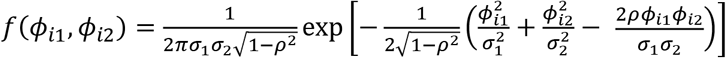. We use Monte Carlo integration to evaluate the log-likelihood function. Let *ϕ*_*i*1*j*_ = *σ*_1_*ξ*_*i*1*j*_ and 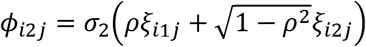, where the *ξ*_*i*1*j*_ and *ξ*_*i*2*j*_ are independently and identically distributed random variables following a standard normal distribution. We have

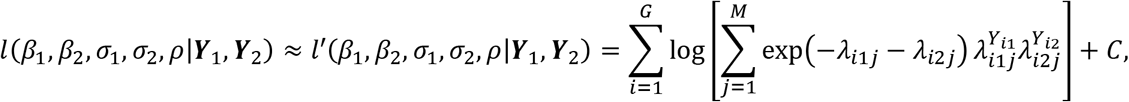

where *λ*_*i*1*j*_ = 2*N*_1_*m_i_* exp(*β*_1_ + *σ*_1_*ξ*_i_1_*j*_) and 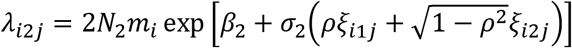. Then, we obtain the MLE of *β*_1_, *β*_2_, *σ*_1_, *σ*_2_, *ρ* through maximization of *l*’(*β*_1_, *β*_2_, *σ*_1_, *σ*_2_, *ρ*|***Y***_1_, ***Y***_2_). Standard error of MLE can be obtained through inversion of the observed Fisher information matrix.

### DNM data and variant annotation

We obtained DNM data from published studies (**Supplementary Table 1**). DNM data for EP from the original release^13^ were not in an editable format and were instead collected from denovo-db^61^. We used ANNOVAR^62^ to annotate all DNMs. Synonymous variants were determined based on the ‘synonymous SNV’ annotation in ANNOVAR; Variants with ‘startloss’, ‘stopgain’, ‘stoploss’, ‘splicing’, ‘frameshift insertion’, ‘frameshift deletion’, or ‘frameshift substitution’ annotations were classified as LoF; Dmis variants were defined as nonsynonymous SNVs predicted to be deleterious by MetaSVM^63^; nonsynonymous SNVs predicted to be tolerable by MetaSVM were classified as Tmis. Other DNMs which did not fall into these categories were removed from the analysis. For each variant class, we estimated the mutability of each gene using a sequence-based mutation model^26^ while adjusting for the sequencing coverage factor based on control trios as previously described^12^ (**Supplementary Table 15**). We included 18,454 autosomal protein-coding genes in our analysis. *TTN* was removed due to its substantially larger size.

### Implementation of mTADA

The software mTADA requires the following parameters as inputs: proportion of risk genes 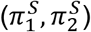, mean relative risks 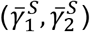, and dispersion parameters 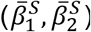 for both disorders. We used extTADA^10^ to estimate these parameters as suggested by the mTADA paper^9^. mTADA reported the estimated proportion of shared risk genes *π*_3_ (posterior mode of *π*_3_) and its corresponding 95% credible interval [LB, UB]. We considered 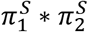 as the expected proportion of shared risk genes, and there is significant genetic sharing between two disorders when 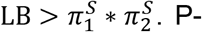. P-value for *π*_3_ was calculated by comparing 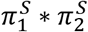 to the posterior distribution of *π*_3_. Number of MCMC chain was set as 2 and number of iterations was set as 10,000.

### Simulation settings

We assessed the performance of EncoreDNM under the mixed-effects Poisson model. We performed simulations for two variant classes: Tmis and LoF variants, which have the largest and the smallest median mutability values across all genes. First, we performed single-trait simulations to assess estimation precision of elevation parameter *β* and dispersion parameter *σ*. We set the true values of *β* to be −0.5, −0.25, and 0, and the true values of *σ* to be 0.5, 0.75, and 1. These values were chosen based on the estimated parameters in real DNM data analyses and ensured simulation settings to be realistic. Next, we performed simulations for cross-trait analysis to assess estimation precision of enrichment correlation *ρ* whose true values were set to be 0, 0.2, 0.4, 0.6, and 0.8. Sample size for each disorder was set to be 5,000. Coverage rate was calculated as the percentage of simulations that the 95% Wald confidence interval covered the true parameter value. Each parameter setting was repeated 100 times.

We also carried out simulations to compare the performance of EncoreDNM and mTADA. Type I error and statistical power for EncoreDNM were calculated as the proportion of simulation repeats that p-value for enrichment correlation *ρ* was smaller than 0.05. and the proportion of simulation repeats that p-value for estimated proportion of shared risk genes *π*_3_ was smaller than 0.05 was used for mTADA. We aggregated all variant classes together, so mutability for each gene was determined as the sum of mutabilities across four variant classes (i.e. LoF, Dmis, Tmis, and synonymous).

First, we simulated DNM data under the mixed-effects Poisson model. To see whether two methods would produce false positive findings, we performed simulations under the null hypothesis that the enrichment correlation *ρ* is zero. We compared two methods under a range of parameter combinations of (*β*, *σ*, *N*) for both disorders: (−0.25, 0.75, 5000) for the baseline setting, (−1, 0.75, 5000) for a setting with small *β*, (−0.25, 0.5, 5000) for a setting with small *σ*, and (−0.25, 0.75, 1000) for a setting with small sample size. We also assessed the statistical power of two methods under the alternative hypothesis. True value of enrichment correlation *ρ* was set to be 0.05, 0.1, 0.15, and 0.2. In the power analysis, parameters (*β*, *σ*, *N*) were fixed at (−0.25, 0.75, 5000) as in the baseline setting when both methods had well-controlled type-I error.

To ensure a fair comparison, we also compared EncoreDNM and mTADA under a multinomial model, which is different from the data generation processes for the two approaches. For each disorder (*k* = 1,2), we randomly selected causal genes of proportion 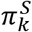. A proportion (i.e., *π*_3_) of causal genes overlap between two disorders. We assumed that the total DNM count to follow a Poisson distribution: 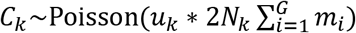, where *u_k_* represents an elevation factor to represent systematic bias in the data. Let ***Y**_k_* denote the vector of DNMs counts in the exome, ***m*** denote the vector of mutability values for all genes, and ***m**_causal,k_* denote the vector of mutability with values set to be 0 for non-causal genes of disorder *k*. We assumed that a proportion *p_k_* of the probands could be attributed to DNMs burden in causal genes, and 1 – *p_k_* of the probands obtained DNMs by chance:

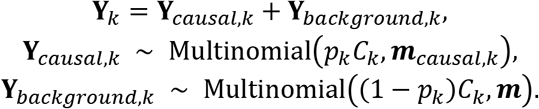

To check whether false positive findings could arise, we performed simulations under the null hypothesis that 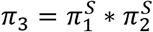 across a range of parameter combinations of (*u*, *p*, *N*, *π^s^*) for both disorders: (0.95, 0.25, 5000, 0.1) for the baseline setting, (0.75, 0.25, 5000, 0.1) for a setting with small *u* (i.e., reduced total mutation count), (0.95, 0.15, 5000, 0.1) for a setting with small *p* (fewer probands explained by DNMs), and (0.95, 0.25, 1000, 0.1) for a setting with smaller sample size. We also assessed the statistical power of two methods under the alternative hypothesis that 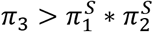. In power analysis, (*u*, *p*, *N*, *π^s^*) were fixed at (0.95, 0.25, 5000, 0.1) as in the baseline setting when type-I error for both methods were well-calibrated.

### Comparison to the fixed-effects Poisson model

For single-trait analysis, the fixed-effects Poisson model assumes that

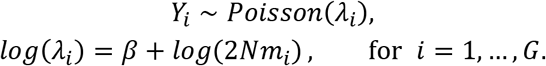

Note that the fixed-effects Poisson model is a special case of our proposed mixed-effects Poisson model when *σ* = 0. We compared the two models using likelihood ratio test. Under the null hypothesis that *σ* = 0, 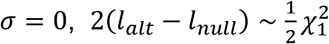 asymptotically, where *l_alt_* and *l_null_* represent the log likelihood of the fitted mixed-effects and fixed-effects Poisson models respectively.

### Recurrent genes and DNMs

We used FUMA^64^ to perform GO enrichment analysis for genes harboring LoF DNMs in multiple disorders. Due to potential sample overlap between the studies of DD^3^ and ID^2^, we excluded ID from the analysis of recurrent DNMs. We calculated the probability of observing two identical DNMs in two disorders using a Monte Carlo simulation method. For each disorder, we simulated exome-wide DNMs profile from a multinomial distribution, where the size was fixed at the observed DNM count and the per-base mutation probability was determined by the tri-nucleotide base context. We repeated the simulation procedure 100,000 times to evaluate the significance of recurrent DNMs. Lollipop plots for recurrent mutations were generated using MutationMapper on the cBio Cancer Genomics Portal^65^.

### Implementation of cross-trait LD score regression

We used cross-trait LDSC^18^ to estimate genetic correlations between disorders. LD scores were computed using European samples from the 1000 Genomes Project Phase 3 data^66^. Only HapMap 3 SNPs were used as observations in the explanatory variable with the --merge-alleles flag. Intercepts were not constrained in the analyses.

### Estimating enrichment correlation in gene sets

Genes with a high/low probability of intolerance to LoF variants (high-pLI/low-pLI) were defined as the 4,614 genes in the upper/lower quartiles of pLI scores^45^. Genes with high/low brain expression (HBE/LBE) were defined as the 4,614 genes in the upper/lower quartiles of expression in the human fetal brain^46^. Genes with high/low heart expression (HHE/LHE) were defined as the 4,614 genes in the upper/lower quartiles of expression in the developing heart of embryonic mouse^67^. Five biological pathways have been reported to be involved in CHD: chromatin remodeling, Notch signaling, cilia function, sarcomere structure and function, and RAS signaling^51^. We extracted 1730 unique genes that belong to these five pathways from the gene ontology database^68^ and referred to the union set as CHD-related genes. We repeated EncoreDNM enrichment correlation analysis in these gene sets. One-sided Kolmogorov-Smirnov test was used to assess the statistical difference between enrichment correlation signal strength in different gene sets.

### URLs

GWAS summary statistics data of ASD, SCZ, and TD were downloaded on the PGC website, https://www.med.unc.edu/pgc/download-results/; Summary statistics of cognitive performance were downloaded on the SSGAC website, https://www.thessgac.org/data; Summary statistics of epilepsy were downloaded on the epiGAD website, http://www.epigad.org/; pLI scores were downloaded from gnomAD v3.1 repository https://gnomad.broadinstitute.org/downloads; mTADA, https://github.com/hoangtn/mTADA; denovo-db, https://denovo-db.gs.washington.edu/denovo-db/; MutationMapper on cBioPortal, https://www.cbioportal.org/mutation_mapper; LDSC, https://github.com/bulik/ldsc.

## Code availability

EncoreDNM software is available at https://github.com/ghm17/EncoreDNM.

## Acknowledgements

LH acknowledges research support from the National Science Foundation of China (Grant No. 12071243) and Shanghai Municipal Science and Technology Major Project (Grant No. 2017SHZDZX01). QL acknowledges research support from the University of Wisconsin-Madison Office of the Chancellor and the Vice Chancellor for Research and Graduate Education with funding from the Wisconsin Alumni Research Foundation and the Waisman Center pilot grant program at University of Wisconsin-Madison. HZ acknowledges research support from the National Institutes of Health (Grant No. R03HD100883)

## Author contribution

H.G., L.H., and Q.L. designed the study.

H.G. performed data analysis and implemented the software.

Y.S. implemented an early version of the method.

S.C.J., X.Z., and B.L. assisted DNM and mutability data preparation.

R.P.L and M.B. advised on disease biology, data interpretation, and genetic issues.

H.Z. and Q.L. advised on statistical issues.

H.G., L.H., and Q.L. wrote the manuscript.

All authors contributed in manuscript editing and approved the manuscript.

