## Supplementary Figures for "Quantifying concordant genetic effects of *de novo* mutations on multiple disorders"

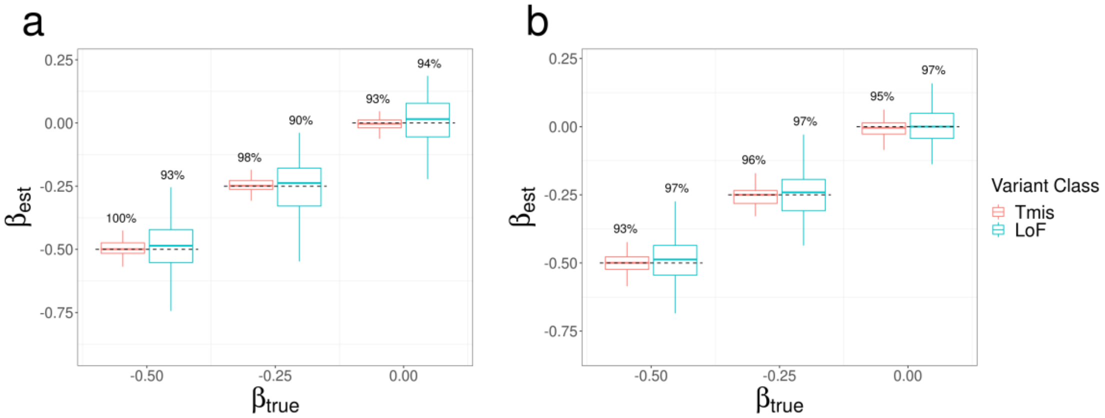


**Supplementary Figure 1. Estimation results of elevation parameter $\beta$ under a mixed-effects Poisson regression model.** Boxplots of estimated value for $\beta$ in single trait analysis with $\sigma_{true}$ value fixed at (a) 0.5 and (b) 1. True parameter values are marked by dashed lines. The number above each box represents the coverage rate of 95% Wald confidence intervals. Each simulation setting was repeated 100 times.

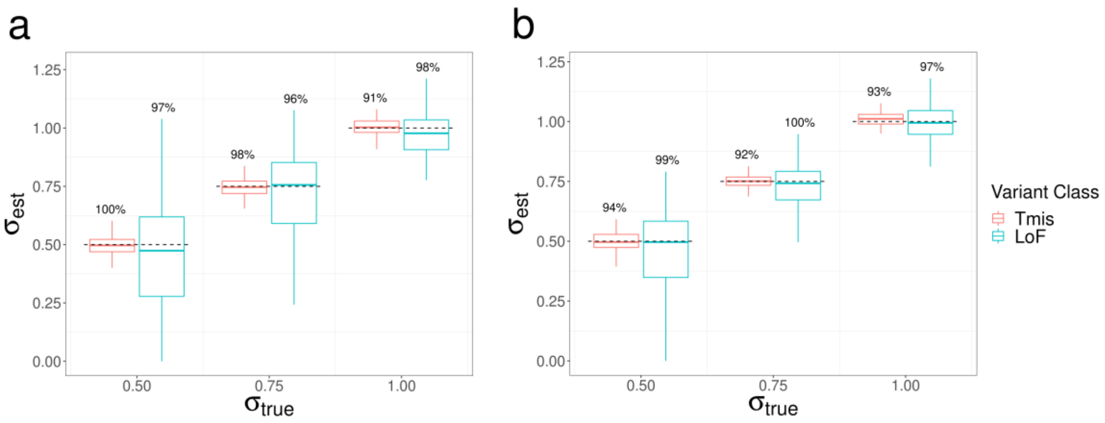


**Supplementary Figure 2. Estimation results of dispersion parameter $\sigma$ under a mixed-effects Poisson regression model.** Boxplots of estimated value for $\sigma$ in single trait analysis with $\beta_{true}$ value fixed at (a) -0.5 and (b) 0. True parameter values are marked by dashed lines. The number above each box represents the coverage rate of 95% Wald confidence intervals. Each simulation setting was repeated 100 times.

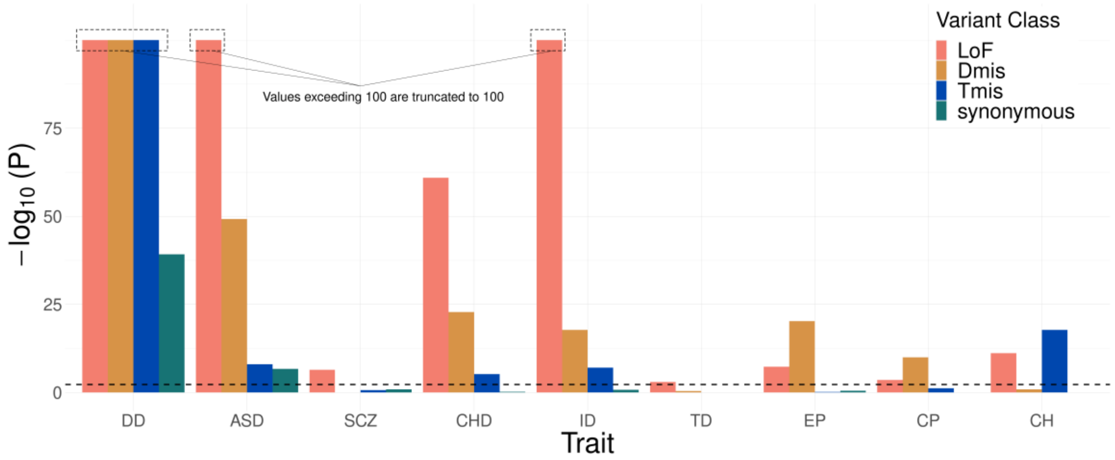


**Supplementary Figure 3. Likelihood ratio test shows significantly improved goodness of fit of the mixed-effects Poisson model compared to a fixed-effects model without the deviation component**. Results with $-\log_{10} P > 100$ are truncated to 100 for visualization purpose.

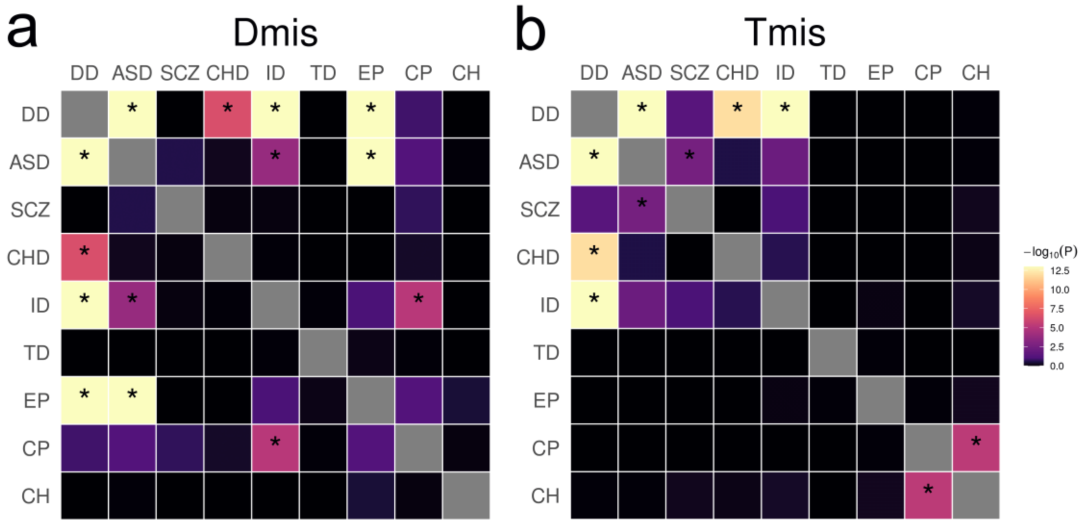


**Supplementary Figure 4. DNM enrichment correlations of nine disorders based on Dmis and Tmis variants.** Significant correlations (FDR<0.05) are marked by asterisks. Results with $-\log_{10} P > 13$ are truncated to 13 for visualization purpose.

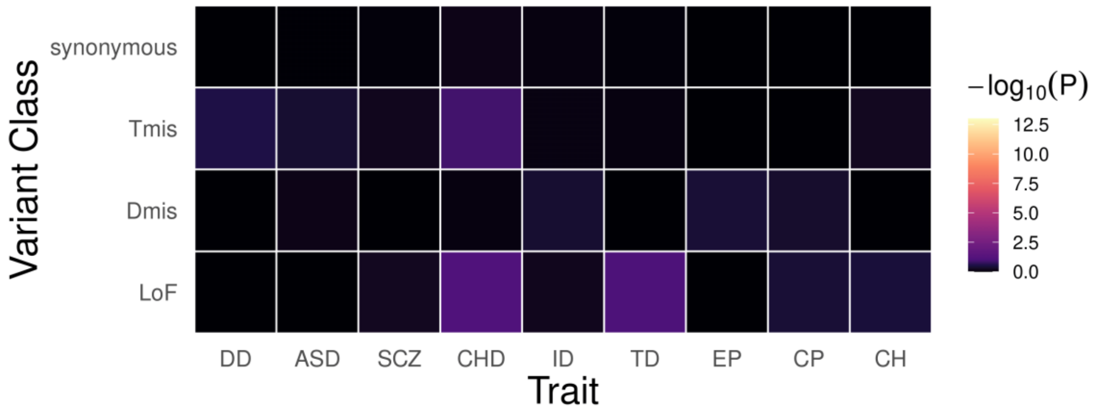


**Supplementary Figure 5. DNM enrichment correlations between nine disorders and controls.**

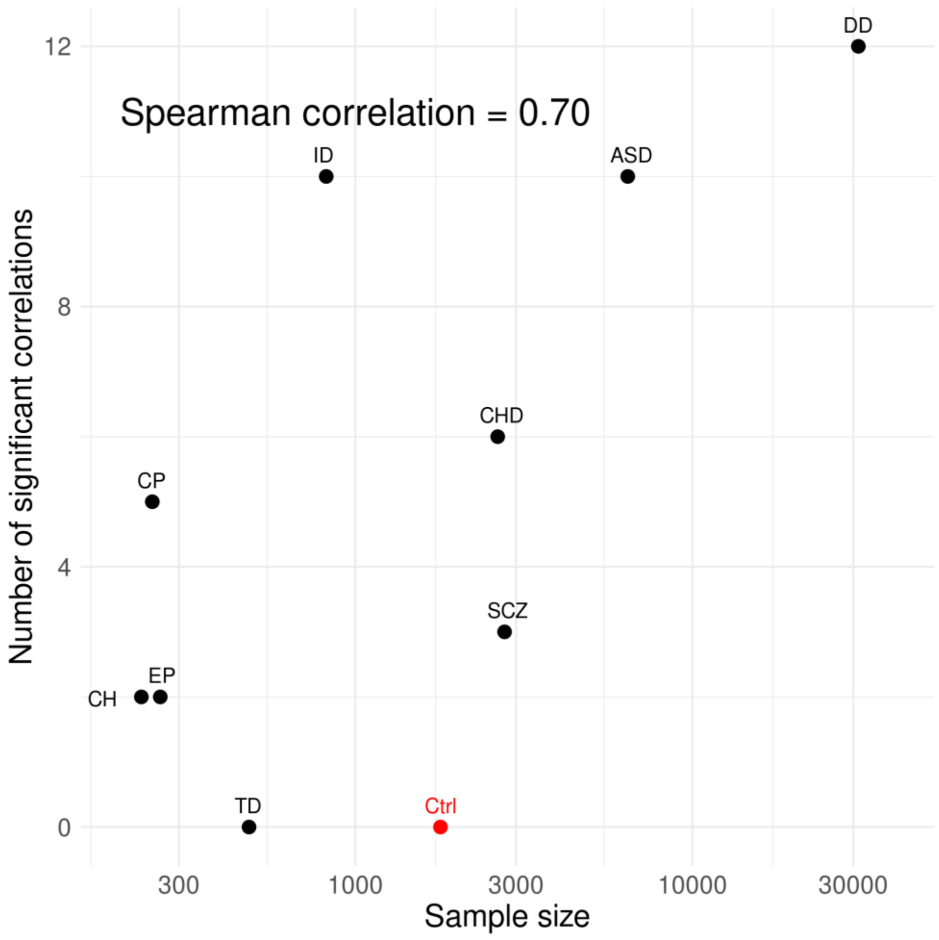


**Supplementary Figure 6. Number of significant correlations identified for each disorder is proportional to its sample size.** X-axis denotes number of trios in each study and is visualized in the log-scale. The data point for controls is a notable outlier and is excluded in the calculation of spearman correlation.

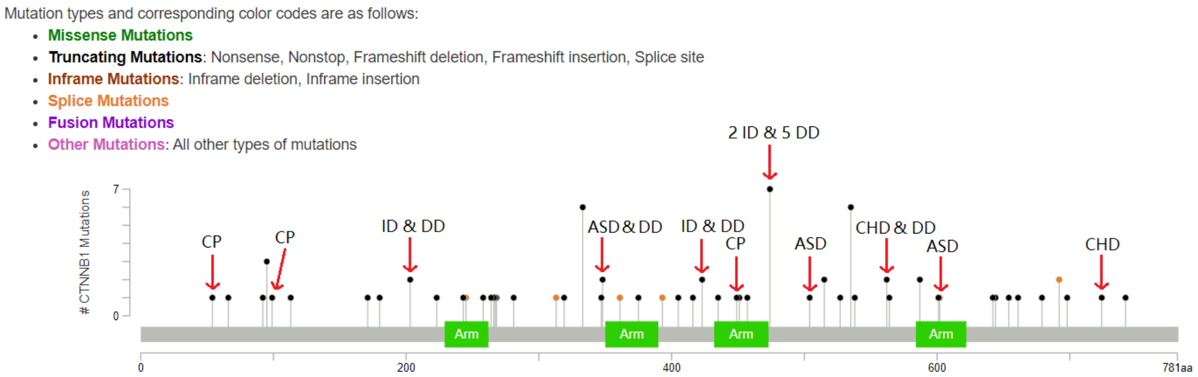


**Supplementary Figure 7. Lollipop plot for LoF DNMs in *CTNNB1*.** Red arrows highlight the DNMs in CP, ID, ASD, and CHD. Other DNMs are from DD probands.

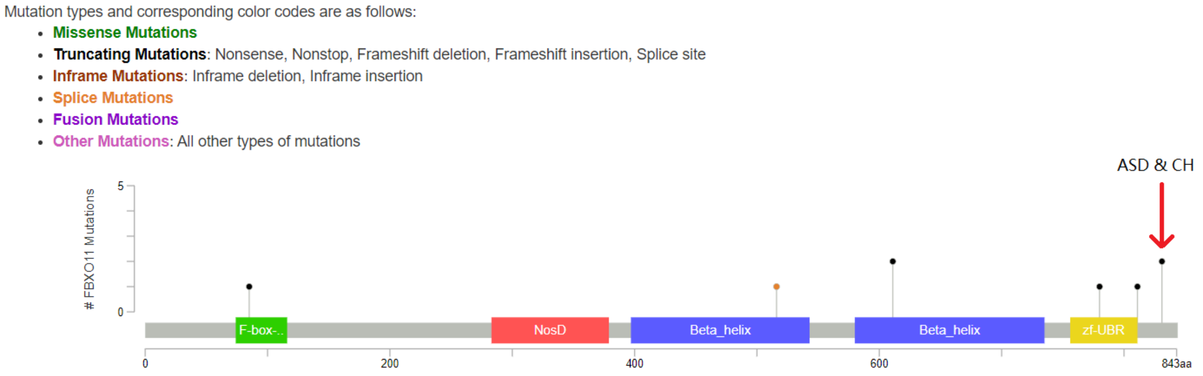


**Supplementary Figure 8. Lollipop plot for LoF DNMs in *FBXO11*.** The red arrow highlights the two identical DNMs in ASD and CH. Other DNMs are from DD probands.

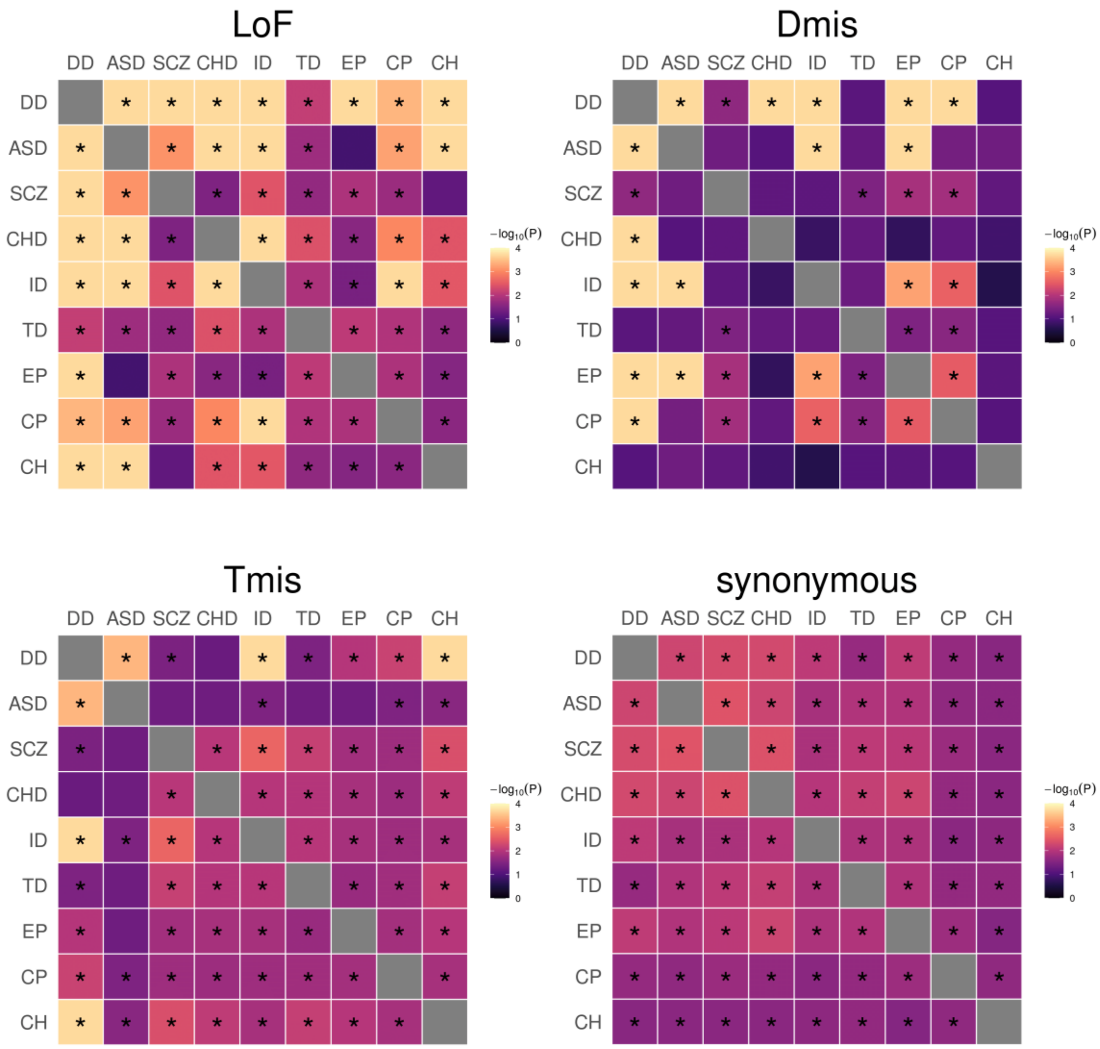


**Supplementary Figure 9. Proportion of shared causal genes in nine disorders estimated for LoF, Dmis, Tmis, and synonymous DNMs using mTADA.** Significant genetic sharings (FDR<0.05) are marked by asterisks.

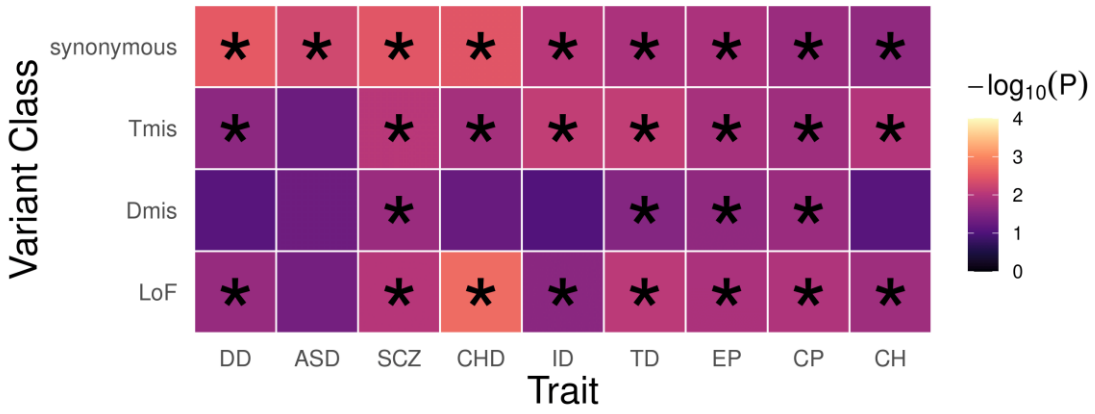


**Supplementary Figure 10. DNM genetic sharing in nine disorders and controls identified by mTADA.**

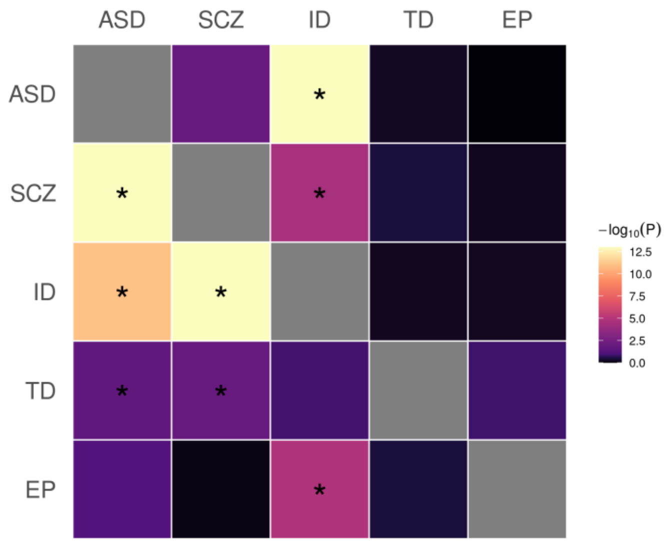


**Supplementary Figure 11. Comparison of GWAS- and DNM-based estimation of genetic sharing among five disorders.** The upper triangle represents enrichment correlations estimated by EncoreDNM. The lower triangle represents genetic correlations estimated from GWAS summary statistics by cross-trait LDSC. We used GWAS on cognitive performance as a proxy for ID. Significant correlations (FDR<0.05) are marked by asterisks. Results with $-\log_{10} P > 13$ were truncated to 13 for visualization purpose.

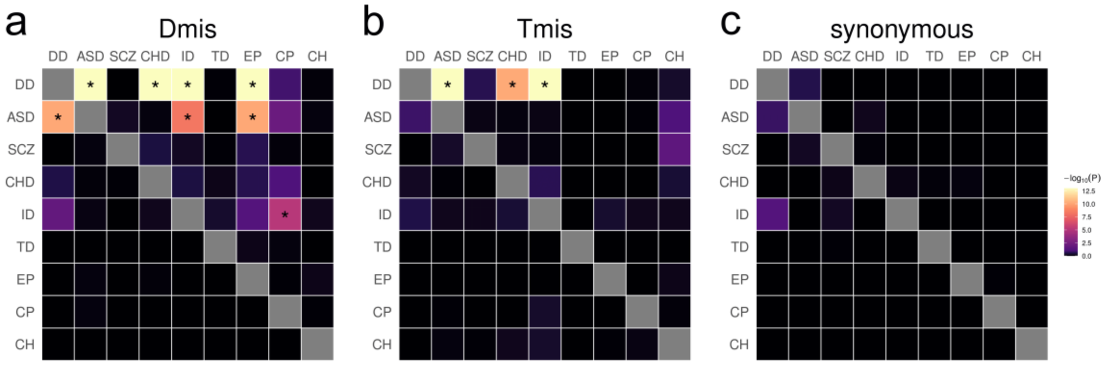


**Supplementary Figure 12. DNM enrichment correlations in high-pLI genes (upper triangle) and low-pLI genes (lower triangle) for Dmis, Tmis, and synonymous variants.** Significant correlations (FDR<0.05) are marked by asterisks. Results with $-\log_{10} P > 13$ are truncated to 13 for visualization purpose.

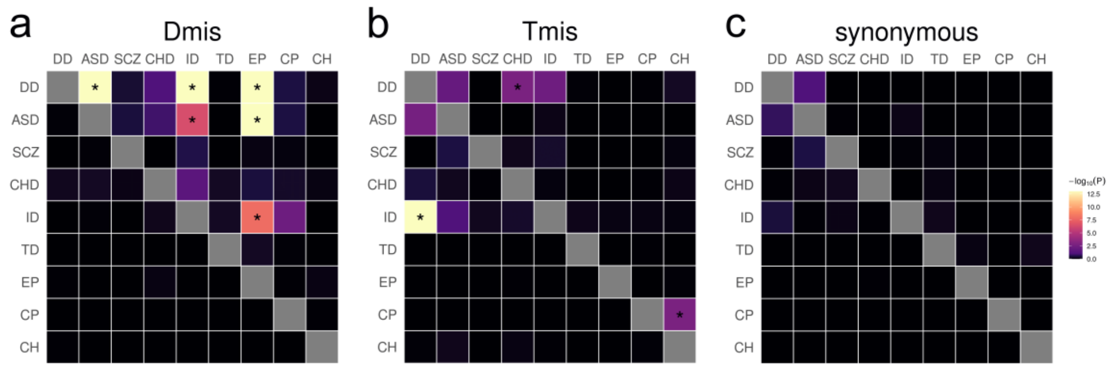


**Supplementary Figure 13. DNM enrichment correlations in HBE genes (upper triangle) and LBE genes (lower triangle) for Dmis, Tmis, and synonymous variants.** Significant correlations (FDR<0.05) are marked by asterisks. Results with $-\log_{10} P > 13$ are truncated to 13 for visualization purpose.

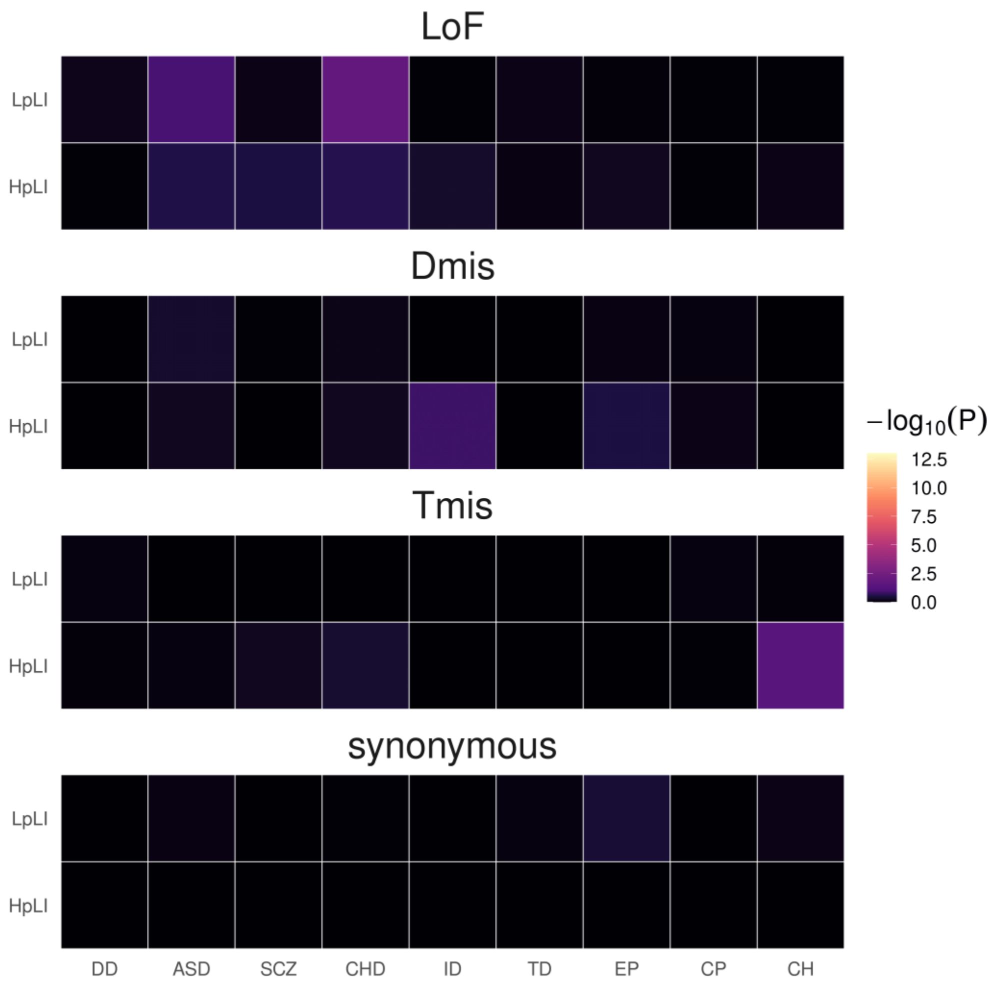


**Supplementary Figure 14. DNM enrichment correlations between nine disorders and controls in high-pLI and low-pLI gene sets.**

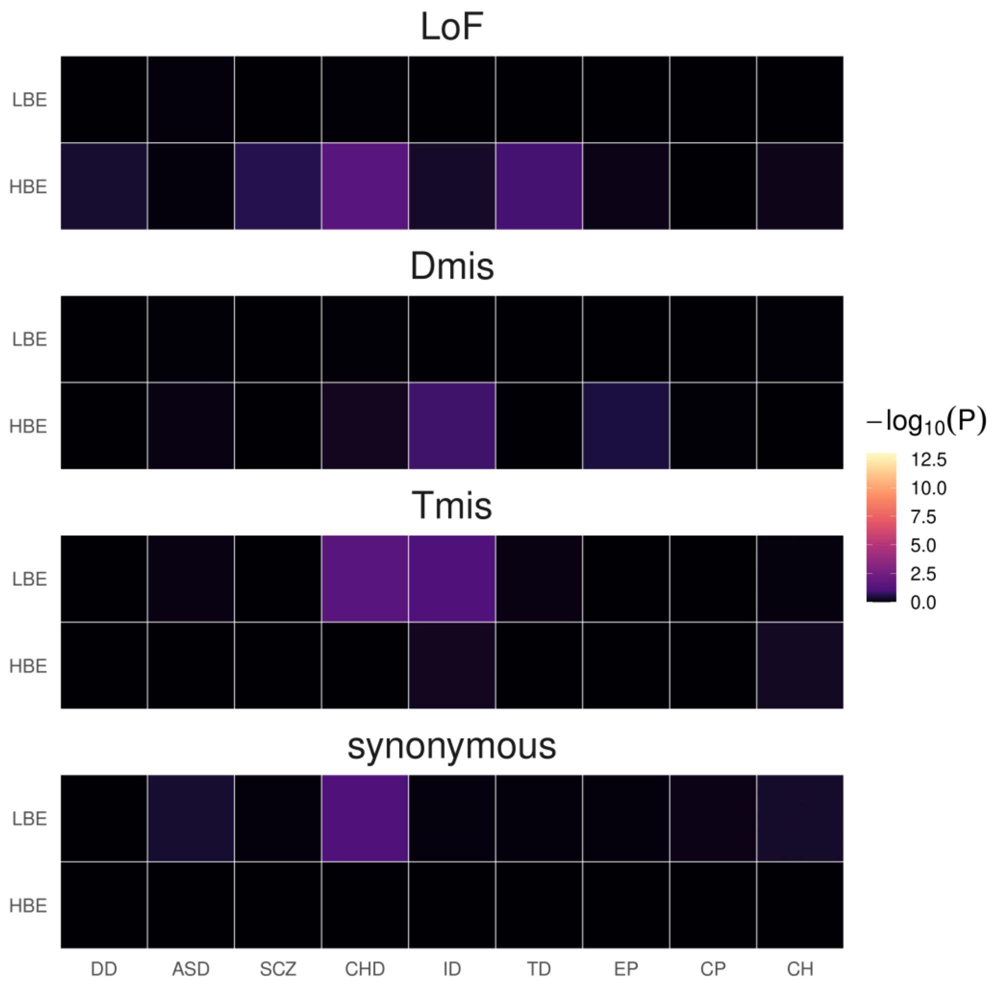


**Supplementary Figure 15. DNM enrichment correlations between nine disorders and controls in HBE and LBE genes.**

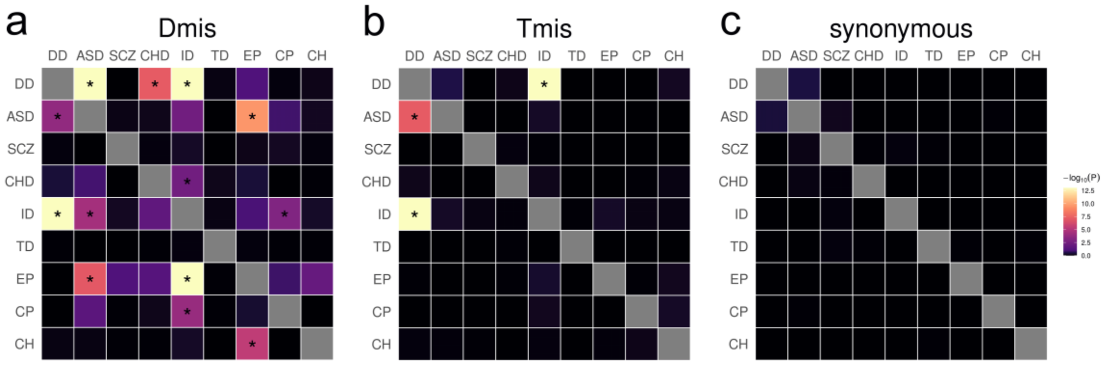


**Supplementary Figure 16. DNM enrichment correlations in HHE genes (upper triangle) and LHE genes (lower triangle) for Dmis, Tmis, and synonymous variants.** Significant correlations (FDR<0.05) are marked by asterisks. Results with $-\log_{10} P > 13$ are truncated to 13 for visualization purpose.

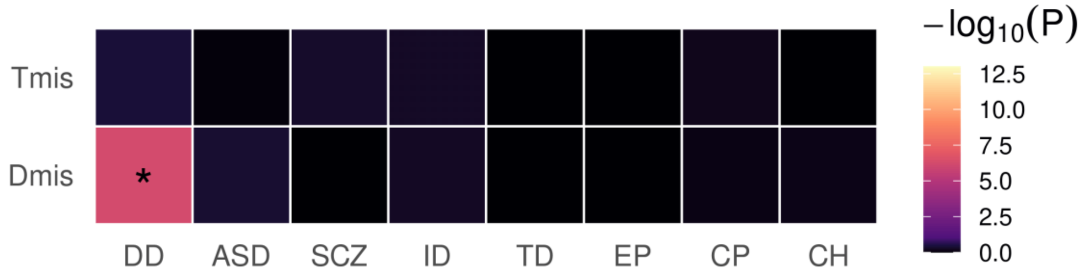


**Supplementary Figure 17. DNM enrichment correlations in CHD-related pathways for Dmis and Tmis variants.** Significant correlations (FDR<0.05) are marked by asterisks.

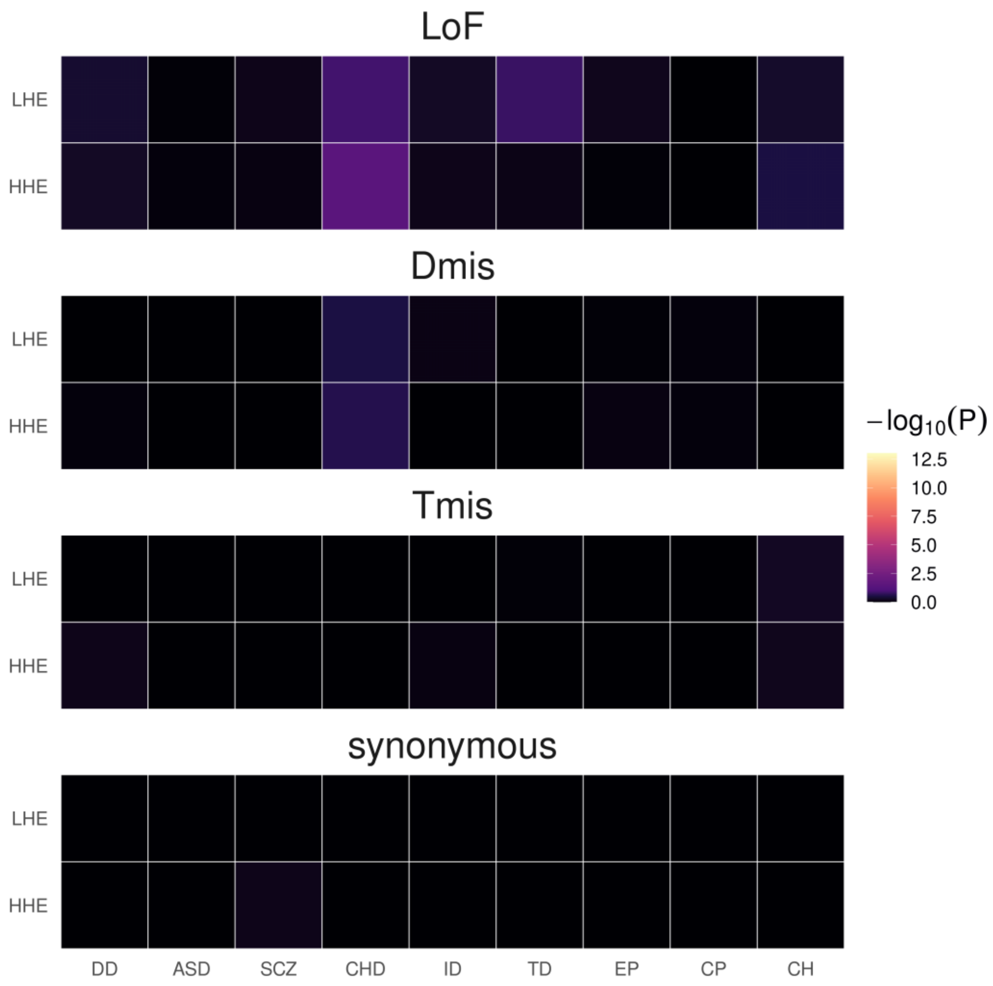


**Supplementary Figure 18. DNM enrichment correlations between nine disorders and controls in HHE and LHE gene sets.**

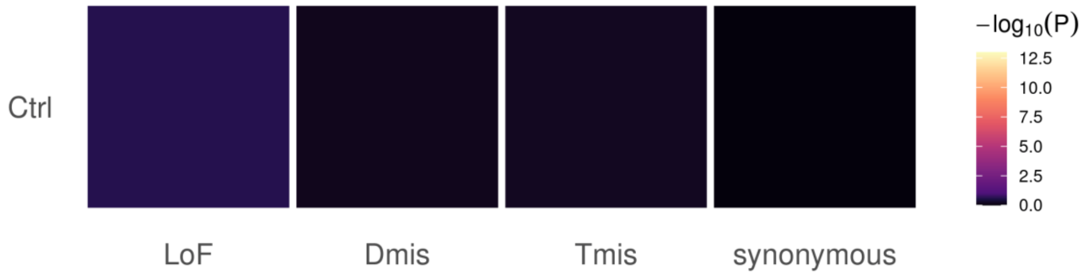


**Supplementary Figure 19. DNM enrichment correlations between CHD and controls in CHD-related pathways.**
